# Single-Cell Super-Resolution Quantification of Oxidative DNA Damage via Aptamer-Assisted DNA-PAINT

**DOI:** 10.1101/2025.08.15.670555

**Authors:** Jingfang Zhao, Li-Sheng Zhang, Yibin Liu, Shuoxing Jiang, Limin Xiang

**Author notes:** Corresponding author., **Materials & Correspondence:** Correspondence and requests for materials should be addressed to L.X.

## Abstract

Accurately quantifying oxidative DNA damage at the single-cell level remains a major challenge due to the limitations of conventional ensemble-based assays, which obscure cell-to-cell variability and lack molecular specificity. To address this, we developed a super-resolution imaging strategy that combines an 8-oxo-dG–specific DNA aptamer with DNA-PAINT, enabling quantitative visualization of 8-oxo-dG lesions with ∼22 nm spatial resolution in individual cells. Our approach reliably detects 8-oxo-dG clusters, distinguishes oxidative damage levels across cells, and exhibits superior specificity and labeling efficiency compared to antibody-based methods. Furthermore, it enables direct evaluation of the repair efficiency of hOGG1 and its catalytic mutants at the single-cell level, overcoming the confounding effects of variable transfection efficiency. This method also serves as a platform for assessing the effects of pharmacological modulators and antioxidants on DNA repair. Looking forward, this strategy can be extended to map other small molecular targets, such as epigenetically modified DNA or RNA bases, with single-cell precision.

## Introduction

Cellular oxidative stress occurs when reactive oxygen species (ROS) overwhelm antioxidant defenses, causing damage to DNA, proteins, and lipids^1-3^. Among oxidative DNA lesions, 8-oxo-7,8-dihydro-2′-deoxyguanosine (8-oxo-dG) is one of the most abundant and mutagenic, capable of mispairing with adenine during replication and inducing GC→TA transversions^4^. Accurate quantification of 8-oxo-dG is therefore essential for assessing oxidative DNA damage and its mutagenic potential, as well as for understanding cellular responses to oxidative stress^5, 6^.

Liquid chromatography–tandem mass spectrometry (LC-MS/MS), enzyme-linked immunosorbent assay (ELISA), and high-throughput sequencing are widely used for detecting 8-oxo-dG^7-13^. LC-MS/MS separates oxidized nucleosides by retention time and quantifies them via mass spectrometry, offering high specificity and sensitivity. ELISA employs antibodies to generate optical signals and is convenient and cost-effective for routine use. More recently, high-throughput sequencing approaches—often coupled with chemical labeling or enrichment strategies—have enabled genome-wide mapping of oxidative DNA lesions, providing base-level resolution and ultra-high sensitivity. However, all these techniques rely on bulk measurements from pooled cellular material, inherently masking cell-to-cell heterogeneity and limiting their utility for analyzing DNA damage and repair dynamics at the single-cell level. In practice, 8-oxo-dG levels can vary substantially among individual cells, as inherent cellular diversity and repeated oxidative challenges drive adaptive responses. This heterogeneity is inaccessible to bulk assays and further hinders evaluation of the DNA repair enzyme hOGG1 and its mutants, since conventional methods cannot readily correlate 8-oxo-dG content with enzyme activity at the single-cell level.

Fluorescence imaging addresses the ensemble limitation by enabling single-cell analysis, typically through measuring the fluorescence intensity of labeled probes^14-17^. However, for small molecules such as 8-oxo-dG, the requirement for a specific recognition molecule poses a significant challenge. Antibodies, while widely used for protein detection, are hindered by their large size, which limits penetration into densely packed chromatin, and are further compromised by epitope masking and nonspecific binding, thereby reducing detection sensitivity. In addition, intensity-based quantification is susceptible to poorly controlled variables—including labeling efficiency, dye concentration, photobleaching, and local environmental effects—ultimately limiting the reproducibility and quantitative accuracy of 8-oxo-dG measurements.

Recent advances in small-molecule–targeting aptamers have introduced probes with low molecular weight, high specificity, programmability, and low immunogenicity, making them well suited for detecting small molecules in cellular environments^18-22^. DNA aptamers that specifically bind 8-oxo-dG have been developed, exhibiting high affinity and single-base discrimination for precise detection^23-25^. As nucleic acids, aptamers are inherently compatible with DNA-PAINT^26-28^ (Point Accumulation for Imaging in Nanoscale Topography), a super-resolution single-molecule localization microscopy (SMLM) technique that quantitatively maps targets with 20–30 nm resolution by counting transient probe– binding events^29^. Combining an 8-oxo-dG–specific aptamer with DNA-PAINT therefore offers a potential approach for high-resolution, molecularly sensitive detection of oxidative DNA damage at the single-cell level, effectively overcoming the limitations of both bulk biochemical assays and intensity-based imaging.

Herein, we developed a strategy that integrates an 8-oxo-dG–specific DNA aptamer with DNA-PAINT super-resolution imaging for quantitative single-cell analysis of oxidative DNA damage. The aptamer provides high specificity and single-base resolution for recognizing 8-oxo-dG, while DNA-PAINT achieves ∼22 nm spatial resolution, enabling direct visualization and counting of individual 8-oxo-dG clusters within the nucleus. This single-molecule–based quantification not only resolves cellular heterogeneity in oxidative damage but also allows functional evaluation of the DNA repair enzyme hOGG1 and its mutants at the single-cell level following transfection. By linking high-specificity molecular recognition with quantitative super-resolution imaging, our approach provides an unprecedented capability to localize, quantify, and compare oxidative DNA lesions with nanometer precision, offering a powerful platform for both mechanistic studies of DNA repair and potential translational applications.

## Results and Discussions

We began by testing two widely used 8-oxo-dG–specific aptamers, OG10 and OGMS1, for targeting 8-oxo-dG in cellular chromatin^23-25^. Their predicted secondary structures are shown in Fig. 1a. To enable DNA-PAINT imaging, a 60-base docking strand (30×TC) was conjugated to the 5′ end of each aptamer (Fig. 1a), allowing reversible hybridization with a fluorophore-labeled imager strand (4×GA–AF647). Live COS-7 cells were exposed to an oxidative solution (KBrO_3_) for one hour, then fixed and incubated with the aptamers to promote specific binding to 8-oxo-dG, followed by replacement of the solution with imaging buffer containing 0.2 nM imager strand. The transient hybridization between docking and imager strands generated bursts of distinct single-molecule fluorescence signals (Fig. 1c, with OG10 aptamer). Epifluorescence and bright-field images confirmed that these signals were localized within the nucleus (Fig. S1a). Approximately 60,000 frames of single-molecule images were acquired, and each detected spot was fitted with a 2D Gaussian function to determine its centroid position. The accumulated localizations yielded the final super-resolution single-molecule localization microscopy (SMLM) image (Fig. 1d, with OG10 aptamer).

**Figure 1.**
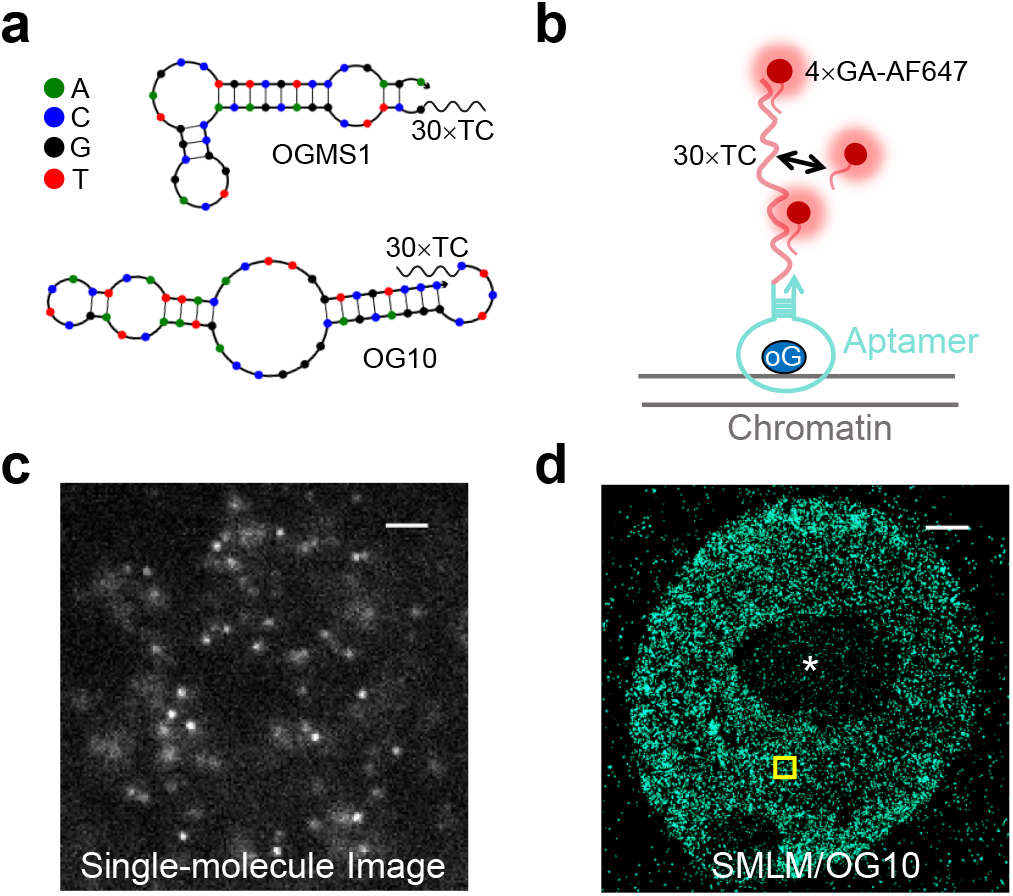
Super-resolution imaging of 8-oxo-dG in cells using aptamer-assisted DNA-PAINT. (a) Predicted secondary structures of the two aptamers used in this study, OG10 and OGMS1. A 30×TC sequence was conjugated to the 5′ end of each aptamer as the docking strand for DNA-PAINT. (b) Schematic illustration of the DNA-PAINT mechanism, where reversible hybridization between the docking strand and fluorophore-labeled imager strand enables detection of 8-oxo-dG in chromatin. (c) Representative single-molecule fluorescence image acquired during DNA-PAINT imaging. (d) Super-resolution single-molecule localization microscopy (SMLM) image reconstructed from millions of localized single-molecule events, revealing numerous 8-oxo-dG clusters within the cell nucleus. Scale bar: 2 μm in **c** and **d**.

Closer examination of the SMLM image revealed numerous 8-oxo-dG clusters (zoom-in in Fig. 2a, corresponding to the yellow box in Fig. 1d), with the exception of nucleolar regions (asterisk in Fig. 1d), known to be enriched in RNA and RNA-associated proteins. A representative cluster is shown in Fig. 2b. Fitting the cross-sectional molecular count profile yielded a full width at half maximum (FWHM) of ∼22 nm, confirming the super-resolution capability of our approach. For quantitative cluster analysis, we applied density-based spatial clustering of applications with noise (DBSCAN). The resulting size distribution (Fig. 2c) ranged from 20 to 150 nm, with a peak at ∼30 nm. The average cluster density within the nucleus was 17.3±2.4 μm^−2^. We also tested OGMS1 and performed control experiments without aptamer and with a 30×TC sequence alone; the quantified cluster densities are summarized in Fig. 2d, with corresponding SMLM images in Fig. S1b. As OG10 yielded higher cluster densities than OGMS1, we selected OG10 for subsequent analyses.

**Figure 2.**
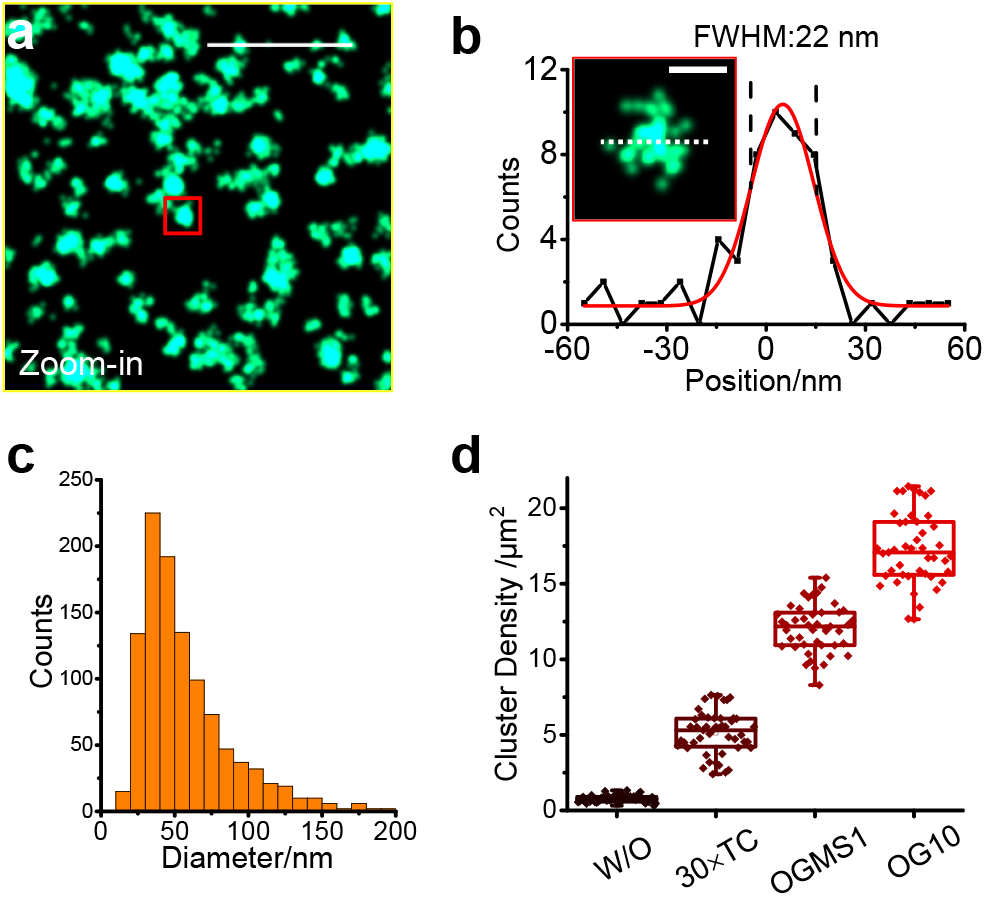
Quantitative analysis of 8-oxo-dG clusters in the cell nucleus. (a) Magnified view of 8-oxo-dG clusters corresponding to the yellow box in Fig. 1d. (b) Cross-sectional profile of single-molecule count from a representative cluster, fitted to determine the full width at half maximum (FWHM), yielding a cluster diameter of ∼22 nm. (c) Statistical distribution of cluster diameters obtained via DBSCAN analysis, with a peak at ∼30 nm. (d) Quantification of cluster density across different binding probes. OG10 demonstrates the highest labeling efficiency among tested aptamers and controls. Scale bar: 500 nm in **a**, 50 nm in **b**. Box plots show median (center line), Box plots show median (center line), interquartile range (box), and outliers (points). >30 regions from ∼10 cells analyzed (**d**).

To compare the performance of OG10 with that of antibodies, we performed two-color SMLM imaging, combining Aptamer-assisted DNA-PAINT for OG10-based detection with immunofluorescence (IF) STORM^30^ (Stochastic Optical Reconstruction Microscopy) imaging of 8-oxo-dG in the green and red channels, respectively. The green channel used 4×GA–ROX for DNA-PAINT, while the red channel employed an AF647-labeled secondary antibody (Fig. 3a). Representative images are shown in Fig. 3b and 3c, where the Aptamer-assisted DNA-PAINT channel via OG10 revealed numerous 8-oxo-dG clusters, whereas the Antibody channel via IF STORM displayed markedly fewer clusters. Epifluorescence images further confirmed the low labeling efficiency of antibody-based detection (Fig. S2). Overlaying the two channels (Fig. 3d) showed that all clusters observed in the Antibody channel were encompassed by those in the OG10 channel. This correspondence was even clearer in the zoom-in view (Fig. 3e), where overlapping clusters appeared white (magenta arrows). Quantitative analysis of cluster density revealed an order-of-magnitude higher density for OG10-based Aptamer-assisted DNA-PAINT compared to antibody-based IF STORM, underscoring the superior labeling efficiency of aptamers for detecting 8-oxo-dG in chromatin.

**Figure 3.**
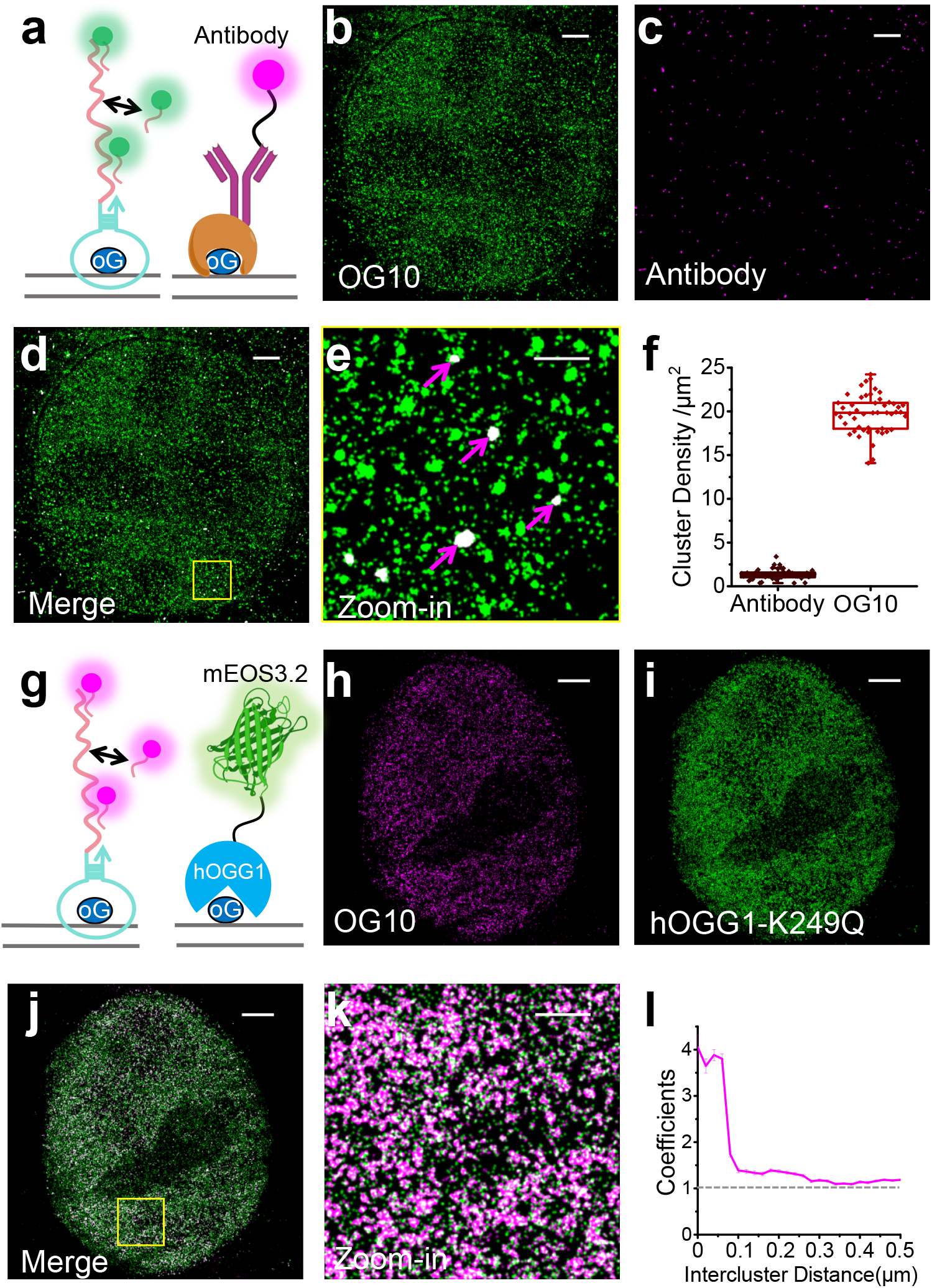
Aptamer-assisted DNA-PAINT imaging of 8-oxo-dG exhibits high labeling efficiency and specificity. (a) Two-color super-resolution imaging of 8-oxo-dG using aptamer-assisted DNA-PAINT (green channel) and antibody-based immunofluorescence (IF) imaging (red channel). (b–d) DNA-PAINT image, IF image, and merged image of 8-oxo-dG from the same cell, showing a higher number of clusters in the DNA-PAINT channel. (e) Zoom-in of (d) reveals that all clusters detected in the antibody channel are encompassed by those in the OG10 aptamer channel. (f) Quantitative comparison of cluster densities between antibody and aptamer channels, demonstrating the superior labeling efficiency of aptamer-assisted DNA-PAINT. (g) Two-color super-resolution imaging of 8-oxo-dG using aptamer-assisted DNA-PAINT (red channel) and PALM imaging of mEOS3.2-tagged hOGG1 K249Q mutant (green channel). (h–j) DNA-PAINT image, PALM image, and merged image from the same cell, showing strong spatial correspondence. (k–l) Zoom-in of (j) and the corresponding cross-correlation analysis confirm high colocalization, highlighting the high specificity of aptamer-assisted DNA-PAINT for 8-oxo-dG detection. Scale bar: 2 μm in **b-d, h-j**, 500 nm in **e** and **k**. Box plots show median (center line), interquartile range (box), and outliers (points). >30 regions from ∼10 cells analyzed (**f**).

To further validate the specificity of our labeling strategy, we performed two-color SMLM imaging, combining aptamer-assisted DNA-PAINT for OG10-based detection of 8-oxo-dG with PALM^31^ (Photoactivated Localization Microscopy) imaging of the mEOS3.2-tagged human 8-oxoguanine DNA N-glycosylase 1 (hOGG1) K249Q mutant in the red and green channels, respectively. The K249Q mutation abolishes hOGG1’s catalytic activity while preserving lesion recognition^32, 33^, resulting in long-lived binding to 8-oxo-dG without cleavage and thus serving as a reliable marker for 8-oxo-dG (Fig. 3g). Representative DNA-PAINT and PALM images (Figs. 3h and 3i) showed comparable cluster densities and spatial distributions. Overlaying the two channels revealed strong spatial correspondence, which was even more evident in the zoom-in view (Fig. 3k). Cross-correlation analysis^34^ (Fig. 3l) further confirmed the high degree of colocalization, providing strong validation for the specificity of our aptamer-based labeling approach.

Having validated our approach for single-cell quantification of 8-oxo-dG, we next examined cells subjected to different levels of oxidative damage. Cells were treated with potassium bromate (KBrO_3_) for varying durations, followed by Aptamer-assisted DNA-PAINT imaging of 8-oxo-dG. Representative images are shown in Fig. 4a, arranged from left to right in order of increasing oxidative damage. A clear rise in cluster density was observed with longer treatment times, as further illustrated in the magnified views and epifluorescence images (Fig. S3). After 2 h of oxidative treatment, 8-oxo-dG labeling became largely continuous, making cluster density quantification by DBSCAN impractical. The statistical analysis of cluster density for quantifiable conditions is summarized in Fig. 4e.

**Figure 4.**
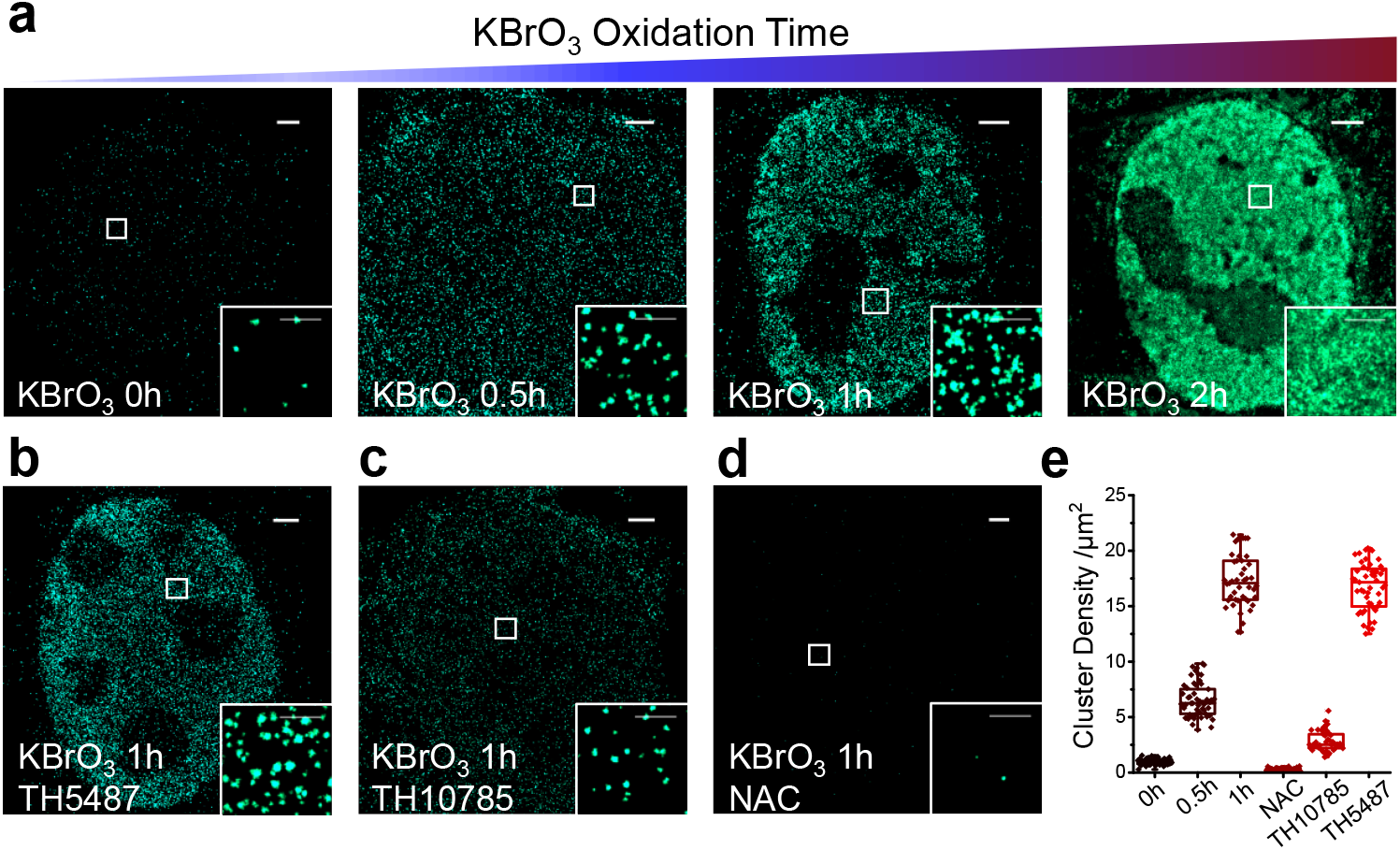
Quantification of 8-oxo-dG content in cells under varying levels of oxidative stress and drug treatment. (a) Aptamer-assisted DNA-PAINT images of cells exposed to oxidative solution (KBrO_3_) for increasing durations, showing a time-dependent increase in 8-oxo-dG cluster density. (b–d) Representative DNA-PAINT images of cells treated with the hOGG1 inhibitor TH5487, activator TH10785, and the antioxidant N-acetylcysteine (NAC), respectively. The resulting cluster densities reflect the expected biological effects of each compound on oxidative DNA damage and repair. (e) Quantitative comparison of 8-oxo-dG cluster densities across different oxidative stress conditions and drug treatments, demonstrating the sensitivity and applicability of the method for evaluating DNA damage and repair responses. Scale bar: 2 μm in **b-d, h-j**, 500 nm in zoom-ins of **b-d, h-j**. Box plots show median (center line), interquartile range (box), and outliers (points). >30 regions from ∼10 cells analyzed (**e**).

The ability of our approach to quantify 8-oxo-dG at the single-cell level also makes it a valuable platform for evaluating drugs that influence oxidative DNA damage repair. Figures 4b and 4c show representative images of cells treated with the hOGG1 inhibitor TH5487 and the hOGG1 activator TH10785. TH10785, a recently reported small molecule, enhances hOGG1’s lyase activity and increases repair efficiency by up to tenfold^35^. In contrast, TH5487 impairs catalytic activity by locking hOGG1 onto chromatin in an inactive, tightly bound state^36^. As expected, inhibitor treatment increased cluster density, whereas activator treatment caused a pronounced decrease, faithfully reflecting their respective effects on repair. We also tested the reductant N-acetylcysteine (NAC), a reactive oxygen species scavenger that slows the generation of 8-oxo-dG. NAC treatment resulted in a dramatic reduction in cluster density, with nearly no detectable clusters (Fig. 4d), indicating strong protection against oxidative chromatin damage. Quantitative cluster density analyses for all drug treatments are summarized in Fig. 4e.

Human 8-oxoguanine DNA N-glycosylase 1 (hOGG1) plays a central role in excising 8-oxo-dG lesions during base excision repair^32^. Figure 5a highlights three critical amino acids within the active site (PDB ID: 1EBM) and their distinct interactions with 8-oxo-dG. His270 forms a hydrogen bond with the 5′ phosphate of 8-oxo-dG, facilitating lesion recognition and discrimination (Fig. 5b); its alanine substitution (H270A) reduces binding affinity by more than 30-fold^37, 38^. Lys249 functions as the catalytic nucleophile, forming a transient covalent bond with the deoxyribose 1′ carbon to enable base excision (Fig. 5b); the K249Q mutant abolishes catalysis while retaining binding, resulting in prolonged lesion engagement^39^. Phe319 stabilizes lesion docking through π–π stacking with the oxidized base (Fig. 5b); the F319A mutation disrupts this interaction, leading to a similarly severe (>30-fold) loss in binding affinity for 8-oxoG-containing DNA and a complete loss of excision activity^38^.

**Figure 5.**
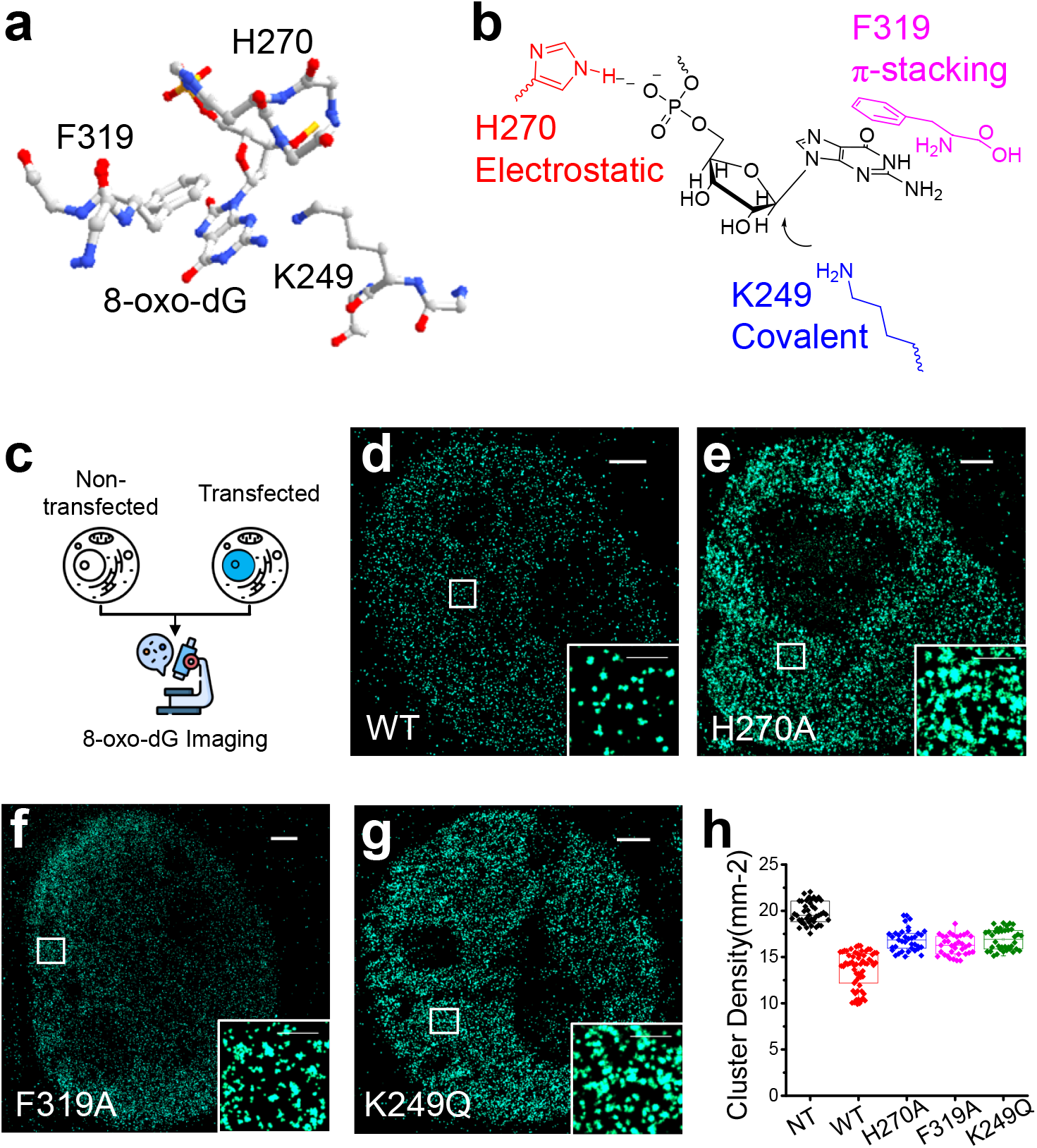
Evaluation of 8-oxo-dG repair efficiency for various hOGG1 variants. (a) Structural interface between hOGG1 and 8-oxo-dG, highlighting three key amino acid residues involved in lesion recognition and excision (PDB ID: 1EBM). (b) Schematic of the distinct interactions mediated by His270, Phe319, and Lys249, each essential for efficient cleavage of 8-oxo-dG. (c) Aptamer-assisted DNA-PAINT imaging of 8-oxo-dG in non-transfected (NT) cells and cells transfected with different hOGG1 variants. (d–g) Representative DNA-PAINT images of cells expressing wild-type hOGG1 and mutant variants (H270A, F319A, and K249Q), showing varying cluster densities. (h) Quantitative comparison of 8-oxo-dG cluster densities across different hOGG1 variants. All mutants exhibit elevated cluster densities compared to wild-type, indicating impaired repair efficiency. Scale bar: 2 μm in **d-g**, 500 nm in zoom-ins of **d-g**. Box plots show median (center line), interquartile range (box), and outliers (points). >30 regions from ∼10 cells analyzed (**h**).

Assessing the impact of hOGG1 mutations on repair efficiency is traditionally achieved through *in vitro* assays such as the comet assay. Evaluating their performance in cells is more challenging, typically requiring silencing of endogenous hOGG1, transfection of mutant constructs, and subsequent bulk quantification of 8-oxo-dG by HPLC or ELISA. Such ensemble measurements cannot resolve cell-to-cell variability—particularly differences in transfection efficiency—thereby masking the true functional effects of individual mutants. Our single-cell approach overcomes this limitation by directly distinguishing non-transfected from successfully transfected cells within the same population (Fig. 5c). Representative 8-oxo-dG images from cells transfected with different variants are shown in Figs. 5d–5g (Fig. S4). Cells expressing wild-type hOGG1 exhibited the lowest cluster density, whereas non-transfected cells showed the highest. All other mutants—including H270A, K249Q, and F319A—displayed intermediate densities, indicating reduced repair efficiency. Statistical comparisons of cluster densities are summarized in Fig. 5h. This ability to selectively quantify 8-oxo-dG in transfected cells provides a powerful and accurate means to evaluate the repair efficiency of hOGG1 mutants under physiologically relevant conditions.

## Conclusions

In summary, we developed a single-cell–resolved, super-resolution imaging strategy that integrates an 8-oxo-dG–specific DNA aptamer with DNA-PAINT to quantitatively assess oxidative DNA damage in the nucleus. This approach offers several key advantages: high specificity for 8-oxo-dG through aptamer-based recognition, nanometer-scale spatial resolution (∼22 nm) through single-molecule localization microscopy, and the ability to resolve cell-to-cell heterogeneity in oxidative damage. Importantly, it enables direct and quantitative evaluation of the repair efficiency of hOGG1 and its mutants within individual transfected cells, bypassing the limitations of ensemble methods. Beyond oxidative DNA damage, this versatile platform provides a generalizable strategy for super-resolution imaging of other small molecular targets, including epigenetically modified DNA and RNA bases, at the single-cell level. Our method therefore represents a powerful and broadly applicable tool for studying nucleic acid modifications^40, 41^ with high spatial precision and molecular sensitivity in complex biological systems.

## Materials and Methods

### Chemical Reagents

KBrO_3_ was purchased from Macklin Biochemical Co., Ltd. TH10785 (CAS No.:1002801-51-5): Obtained from TargetMol, used as an activator. TH5487(CAS No.:2304947-71-3), an inhibitor of hOGG1, Purchased from Shanghai Yuanye Bio-Technology Co., Ltd.,. N-Acetyl-L-Cysteine(NAC)(CAS No.:616-91-1) was ordered from Macklin Biochemical Co., Ltd. STORM Imaging Buffer: Prepared with 100 mM Tris–HCl (pH 8.0), 20 mM NaCl, 100 mM cysteamine, and 10% glucose. An oxygen-scavenging system comprising 60 mg/mL glucose oxidase and 6 mg/mL catalase (both from Sigma-Aldrich) was added to minimize photobleaching during fixed-cell STORM imaging experiments. Blocking Buffer (BB) in immunolabeling: 3% BSA and 0.5% Triton X-100 in PBS. Stored at 4 °C. Washing Buffer (WB) in immunolabeling: 0.2% BSA and 0.1% Triton X-100 in PBS. Stored at 4 °C. Aptamer Binding Buffer: 5 mM MgCl_2_, 0.1 mg/ml Salmon dsDNA (Thermofisher), 1 mg/ml dextran sulphate sodium, 0.1 mg/ml RNase A (Thermofisher), and 0.5% Triton-X in 1⨯PBS. Aptamer Imaging Buffer: 5 mM MgCl_2_, 0.1 mg/ml Salmon dsDNA, and 0.1 mg/ml dextran sulphate sodium in 1⨯PBS. All DNA sequences in Table S1 were synthesized by Sangon Biotech (Shanghai, China).

### Plasmid Construction

The mEos3.2-C1 plasmid (Addgene #54550) was a gift from Michael Davidson and Tao Xu, was used as a backbone to construct hOGG1-mEos3.2 by fusing the hOGG1 sequence (Addgene #18709, provided by David Sidransky) to the N-terminus of mEos3.2 at the SacI and BamH1 restriction site. Mutations K249Q, H270A, and F319A within the hOGG1 coding sequence were introduced using the Phusion Site-Directed Mutagenesis Kit (ThermoFisher Scientific) based on the full-length mEos3.2-tagged hOGG1. Detailed information on all plasmids is summarized in Table S1, and all sequences were verified through Sanger sequencing.

### Coverslips Preparations

Glass coverslips (18 mm in diameter) were first cleaned by immersion in freshly prepared piranha solution (3:1 mixture of sulfuric acid and hydrogen peroxide) heated for 15–20 minutes. After cooling, the coverslips were thoroughly rinsed with Milli-Q water (18.4 MΩ·cm) and dried under a stream of nitrogen gas. The cleaned coverslips were then placed into a 12-well plate and sterilized by UV irradiation for 15 minutes in a biosafety cabinet.

### Cell Culturing and Transfection

COS-7 cells (Cell Bank, GNO 2) were cultured in Dulbecco’s Modified Eagle’s Medium (DMEM) supplemented with 10% fetal bovine serum (FBS), 1% GlutaMAX and 1% penicillin-streptomycin in a humidified incubator at 37°C with 5% CO_2_. For transfected cells, approximately 48–72 hours before imaging, cells were electroporated using a Bio-Rad electroporation system (100 V, 20 ms, 0.2 cm cuvette) in electroporation buffer containing ∼10^5^ cells and 5 μg plasmid DNA per sample. The cell suspension was then plated onto the prepared coverslips with 1 mL of freshly warmed culture medium and mixed gently.

### Oxidative Treatment, Drug Treatment, and Cell Fixation

To induce oxidative DNA damage, cells were treated with 20 mM KBrO_3_ in serum-free DMEM at 37 °C for defined durations (0, 0.5, 1, or 2 h) to promote 8-oxo-dG formation. After treatment, cells were incubated in 1 mL of fresh complete medium for 6 h. For drug treatment, either 10 μM TH10785 (activator), 10 μM TH5487 (inhibitor), 5 mM N-acetylcysteine (NAC, reductant), or an equivalent volume of DMSO (vehicle control) was added to the medium during this 6-hour incubation. Cells were then fixed with freshly prepared 4% (w/v) paraformaldehyde (PFA) in PBS for 20 minutes at room temperature.

### Aptamer Labeling

Following oxidative treatment and fixation, cells were incubated with 100 nM annealed aptamer—30×TC-OG10, 30×TC-OGMS1, or 30×TC alone—in 500 μL of aptamer binding buffer at 4 °C for 1 hour. Aptamers were pre-annealed by heating to 85 °C for 5 minutes in binding buffer, followed by gradual cooling to room temperature. After incubation, unbound aptamers were removed by washing with ice-cold aptamer binding buffer (500 μL for 1 minute).

### DNA-PAINT Imaging

DNA-PAINT imaging was performed by incubating samples with 0.2 nM AF647-conjugated imager strand (5′-GAGAGAGA-3′-AF647) or ROX-conjugated imager strand (5′-GAGAGAGA-3′-ROX) in aptamer imaging buffer. To distinguish transfected from non-transfected cells, epifluorescence images of mEOS3.2 were acquired prior to DNA-PAINT imaging. Imaging was conducted using a custom-built microscope equipped with a 642 nm laser operating at a power density of ∼0.1 kW/cm^2^. Single-molecule fluorescence images were acquired at a frame rate of ∼125 Hz with an effective exposure time of 2 ms per frame. Emission signals were filtered using a longpass filter (ET655lp, Chroma) and a bandpass filter (ET705/100m, Chroma).

### Two-color SMLM Imaging of Aptamer-assisted DNA-PAINT and K249Q hOGG1 mutant-mEOS3.2

K249Q-mEOS3.2–transfected cells were subjected to oxidative treatment and fixation as described above. Aptamer labeling of 8-oxo-dG was performed using 100 nM pre-annealed aptamer in binding buffer at 4 °C for 1 h, following the previously described protocol. For two-color imaging, conventional epifluorescence images of the AF647 signal (red channel) were first acquired to confirm aptamer binding. Aptamer-assisted DNA-PAINT imaging was then conducted using 0.2 nM AF647-labeled imager strand with 642 nm laser excitation (∼0.1 kW/cm^2^) at a frame rate of 125 Hz and an effective exposure time of 2 ms. Fluorescence signals were filtered through a longpass filter (ET655lp, Chroma) and a bandpass filter (ET705/100m, Chroma). To identify transfected cells, epifluorescence imaging of mEOS3.2 (green channel) was performed using 488 nm illumination (∼0.5–1 kW/cm^2^) with 100 ms exposure time. Subsequently, SMLM imaging of the K249Q-mEOS3.2 signal was carried out using 405 nm photoactivation (∼0.5– 1 kW/cm^2^) and 561 nm excitation (∼1–2 kW/cm^2^) at 145 Hz and an effective exposure time of 2 ms. Emission was filtered using a longpass filter (ET575lp, Chroma) and a bandpass filter (ET605/70m, Chroma).

### Two-Color SMLM Imaging of Aptamer-Assisted DNA-PAINT and Immunofluorescence-Based STORM

Following oxidative treatment and fixation, cells were washed three times with DPBS. For immunolabeling, cells were incubated overnight at 4 °C in 1 mL blocking buffer. Primary antibody staining was performed using anti-8-oxo-dG antibody (1:200 in blocking buffer) for 1 h at room temperature (RT), followed by three 10-min washes with washing buffer. Cells were then incubated with AF647-conjugated secondary antibody (1:400 in blocking buffer) for 1 h at RT, followed by a final 10-min wash. Aptamer labeling (100 nM) was subsequently carried out as described previously.

To minimize spectral crosstalk, imaging was performed in the following sequence:

1. Conventional epifluorescence of antibody-labeled 8-oxo-dG (642 nm excitation, 0.5 mW, 100 ms exposure, ET655lp and ET705/100m filters).
2. STORM imaging of antibody-labeled 8-oxo-dG (642 nm, ∼0.1 kW/cm^2^, 125 Hz).
3. Conventional epifluorescence imaging of aptamer signal (560 nm excitation, ∼0.5 mW/cm^2^, 100 ms exposure, ET575lp and ET605/70m filters).
4. Aptamer-assisted DNA-PAINT super-resolution imaging using 0.2 nM ROX-imager strand (560 nm excitation, 125 Hz, 2 ms exposure time).

Channel alignment was verified using 100 nm TetraSpeck beads. All aptamer-related steps were performed at 4 °C to maintain complex stability.

### Optical setup

All super-resolution imaging experiments were conducted using a custom-built system based on an Olympus IX-83 inverted fluorescence microscope. The illumination system integrated five laser lines: 405 nm (CNI MDL-III-405, 500 mW), 488 nm (Coherent OBIS 488 LX, 150 mW), 532 nm (CNI OEM-U-532, 500 mW), 560 nm (MPB Communications, 500 mW), and 642 nm (MPB Communications, 500 mW). Laser beams were focused at the back focal plane of a high-NA oil-immersion objective (Olympus UPLXAPO 100×, NA 1.45), using slightly subcritical angle illumination to achieve ∼2–3 μm penetration depth while maintaining TIRF-like sectioning. Temporal control of the 560 nm and 642 nm laser outputs was achieved via an acousto-optic tunable filter (AOTF, Gooch & Housego) synchronized with camera acquisition using a National Instruments PCI-6733 I/O board. This setup coordinated AOTF modulation with TTL signals from a sCMOS camera (Teledyne Photometrics Prime 95B) and laser triggering pulses, enabling precise 2 ms illumination cycles optimized for single-molecule localization.

### Cluster Density Analysis of 8-oxodG in Fixed Cells

Single-molecule localization clustering was analyzed using density-based spatial clustering of applications with noise (DBSCAN). Two parameters were defined: the maximum distance between neighboring points (ε) and the minimum number of localizations required to form a cluster (N□i□). For 8-oxo-dG analysis in nuclei, a neighborhood radius of 30 nm was used, and clusters containing fewer than 12 localizations within this radius were excluded from analysis.

## Acknowledgments

L.X. and S. J. acknowledges financial supports from National Key R&D Program of China (2022YFA1305400), National Natural Science Foundation of China (22274122, 22104113), Fundamental Research Funds for the Central Universities interdisciplinary (2042023kf1012), Hubei Provincial Natural Science Foundation of China 2025AFA021, and Innovative Talents Foundation from Renmin Hospital of Wuhan University (JCRCFZ-2022-010). S. J. acknowledges financial support from National Natural Science Foundation of China (22302092).

## Author contributions

Conceptualization: L. X., J. Z. Methodology: L. X., J. Z., B. L. Investigation: J. Z. Visualization: L. X., J. Z. Project administration: L. X. Supervision: L. X., S. J. Writing-original draft: L. X., J. Z. Writing-review & editing: L. X., J. Z., B. L., L. Z.

## Materials & Correspondence

Correspondence and requests for materials should be addressed to L.X.

## Supplemental information

Supplementary Information: Figs. S1-S4, Table S1

