## Supplementary Figures for "Single-Cell Super-Resolution Quantification of Oxidative DNA Damage via Aptamer-Assisted DNA-PAINT"

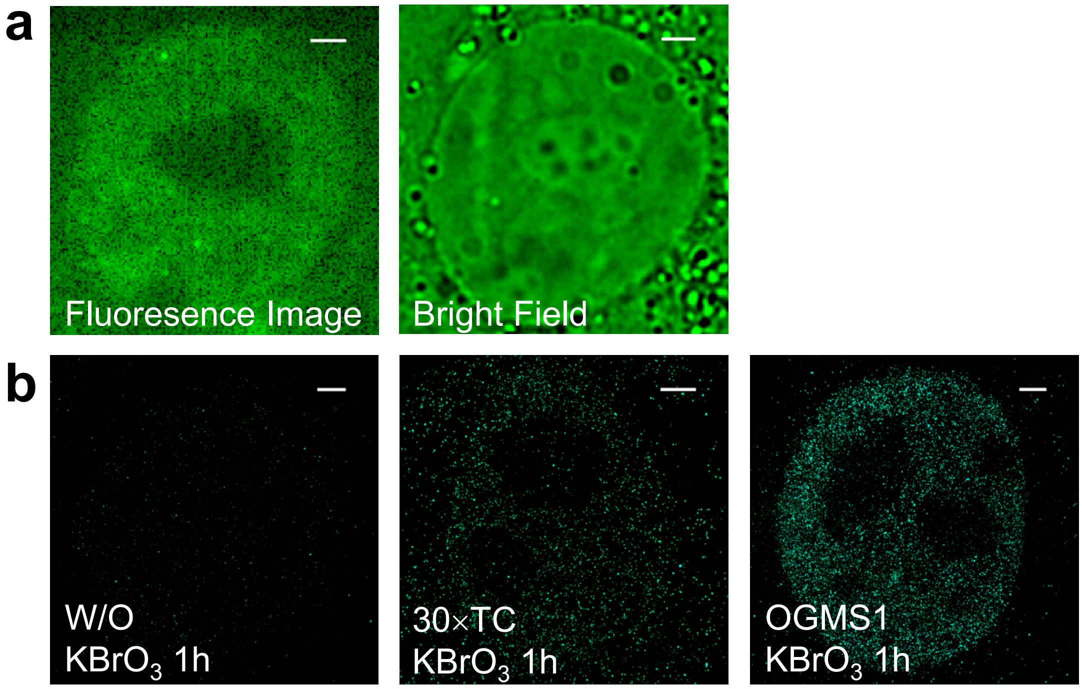


**Figure S1. Aptamer-assisted DNA-PAINT imaging of 8-oxo-dG in fixed cell nuclei.**

(a) Fluorescence image of 8-oxo-dG labeled with the OG10 aptamer.

(b) Brightfield image of fixed cell nuclei.

(c) DNA-PAINT image of 8-oxo-dG in untreated cells, serving as a negative control.

(d) DNA-PAINT image of 8-oxo-dG using a 30×TC sequence without an aptamer, showing minimal labeling.

(e) DNA-PAINT image of 8-oxo-dG labeled with the OGMS1 aptamer..

Scale bar: 2 μm.

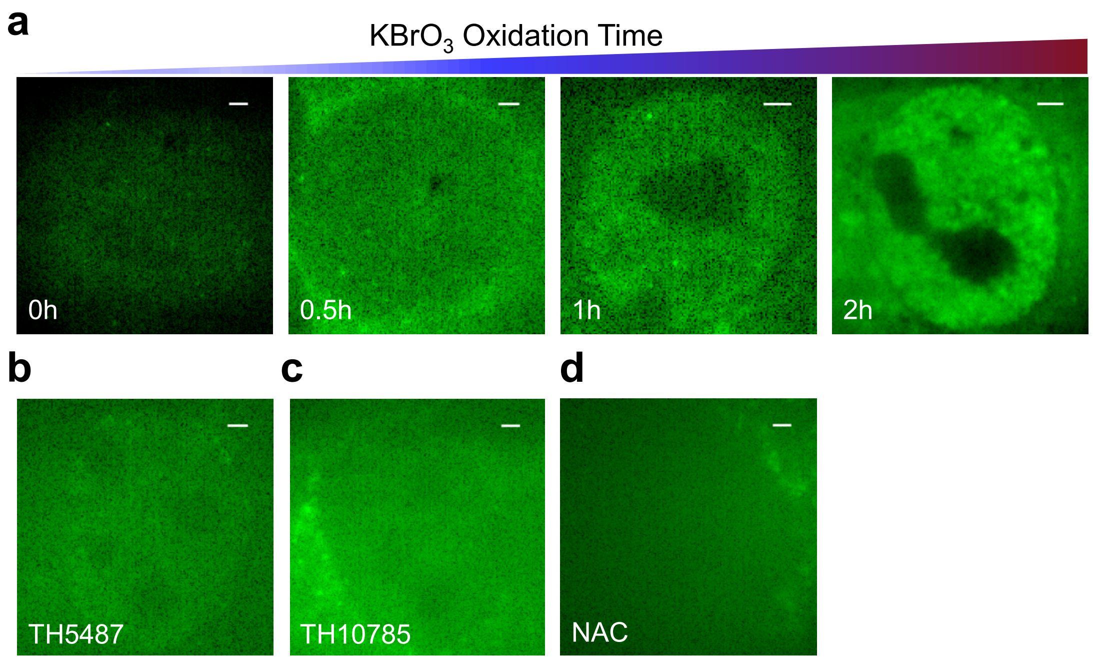


**Figure S2. Fluorescence imaging of 8-oxo-dG labeled with OG10 under varying oxidative stress and pharmacological treatments.**

(a) Fluorescence images of 8-oxo-dG in fixed cell nuclei following potassium bromate ($KBrO_3$) treatment for 0 h (control), 0.5 h, 1 h, and 2 h, showing time-dependent increasing of fluorescence intensity.

(b) Fluorescence image of 8-oxo-dG in cells treated with the hOGG1 inhibitor TH5487.

(c) Fluorescence image of 8-oxo-dG in cells treated with the hOGG1 activator TH10785.

(d) Fluorescence image of 8-oxo-dG in cells treated with the antioxidant N-acetylcysteine (NAC)

Scale bar: 2 μm.

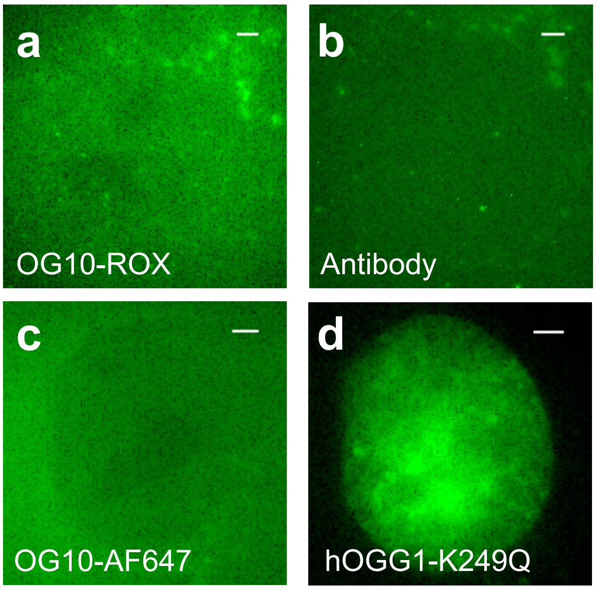


**Figure S3. Fluorescence imaging of 8-oxo-dG and hOGG1 K249Q mutant in fixed cell nuclei.**

(a) Fluorescence image of 8-oxo-dG labeled with OG10-ROX (green channel).

(b) Immunofluorescence image of 8-oxo-dG labeled with anti-8-oxoG antibody and AF647-labeled secondary antibody (red channel).

(c) Fluorescence image of 8-oxo-dG labeled with OG10-AF647 (red channel).

(d) Fluorescence image of hOGG1 K249Q mutant expression in cell nuclei (green channel).

Scale bar: 2 μm.

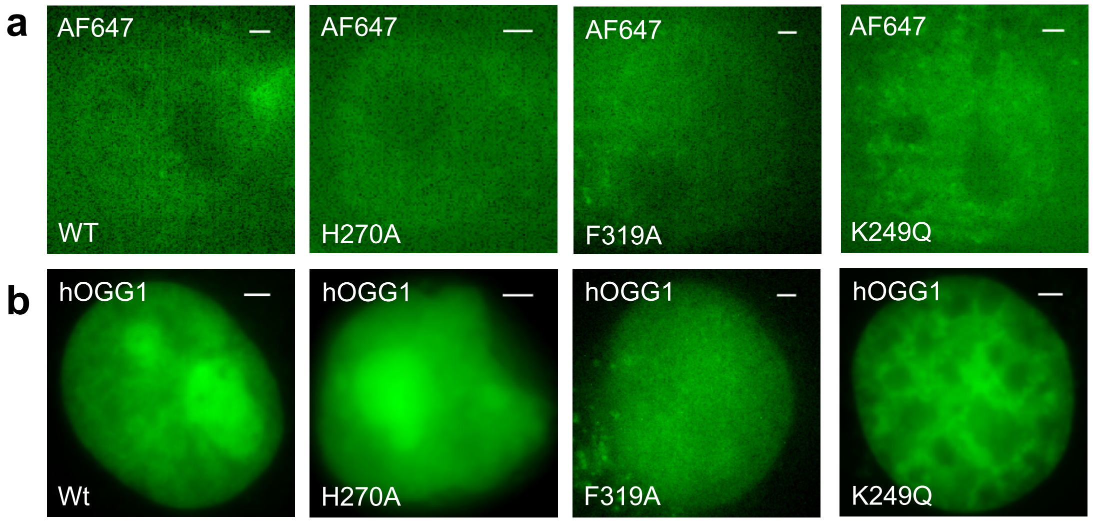


**Figure S4. Fluorescence imaging of 8-oxo-dG and hOGG1 variants in fixed cell nuclei.**

(a) Fluorescence images of 8-oxo-dG labeled with OG10-AF647 in nuclei of cells transfected with wild-type hOGG1 or the H270A, F319A, and K249Q mutants.

(b) Corresponding epifluorescence images showing the expression of transfected hOGG1 variants (wild-type, H270A, F319A, and K249Q).

Scale bar: 2 μm.

**Table S1. The sequences of the all the plasmids and aptamers used in this work**

| Name | Mutated amino acids in proteins and aptamer sequences in DNA were underlined. Docking sequence was Italic. |
|---|---|
| hOGG1 | MPARALLPRRMGHRTLASTPALWASIPCPRSELRLDLVLPSG QSFRWREQSPAHWSGVLADQVWTLTQTEEQLHCTVYRGDK SQASRPTPDELEAVRKYFQLDVTLAQLYHHWGSVDSHFQEV AQKFQGVRLLRQDPIECLFSFICSSNNNIARITGMVERLCQAF GPRLIQLDDVTYHGFPSLQALAGPEVEAHLRKLGLGYRARY VSASARAILEEQGGLAWLQQLRESSYEEAHKALCILPGVGTK VADCICLMALDKPQAVPVDVHMWHIAQRDYSWHPTTSQAK GPSPQTNKELGNFFRSLWGPYAGWAQAVLFSADLRQSRHA QEPPAKRRKGSKGPEG |
| mEOS3.2 | MSAIKPDMKIKLRMEGNVNGHHFVIDGDGTGKPFEGKQSM |

| | |
|---|---|
| | DLEVKEGGPLPFAFDILTTAFHYGNRVFAKYPDNIQDYFKQS<br>FPKGYSWERSLTFEDGGICNARNDITMEGDTFYNKVRFYGT<br>NFPANGPVMQKKTLKWEPSTEKMYVRDGVLTGDIEMALLL<br>EGNAHYRCDFRTTYKAKEKGVKLPGAHFVDHCIEILSHDKD<br>YNKVKLYEHAVAHSGLPDNARR |
| hOGG1-mEOS3.2 (611 AAs) | MAQVQLQVDMPARALLPRRMGHRTLASTPALWASIPCPRSE<br>LRLDLVLPSGQSFRWREQSPAHWSGVLADQVWTLTQTEEQL<br>HCTVYRGDKSQASRPTPDELEAVRKYFQLDVTLAQLYHHW<br>GSVDSHFQEVAQKFQGVRLLRQDPIECLFSFICSSNNNIARIT<br>GMVERLCQAFGPRLIQLDDVTYHGFPSLQALAGPEVEAHLR<br>KLGLGYRARYVSASARAILEEQGGLAWLQQLRESSYEEAHK<br>ALCILPGVGTKVADCICLMALDKPQAVPVDVHMWHIAQRD<br>YSWHPTTSQAKGPSPQTNKELGNFFRSLWGPYAGWAQAVL<br>FSADLRQSRHAQEPPAKRRKGSKGPEGGPVATMSAIKPDMK<br>IKLRMEGNVNGHHFVIDGDGTGKPFEGKQSMDLEVKEGGPL<br>PFAFDILTTAFHYGNRVFAKYPDNIQDYFKQSFPKGYSWERS<br>LTFEDGGICNARNDITMEGDTFYNKVRFYGTNFPANGPVMQ<br>KKTLKWEPSTEKMYVRDGVLTGDIEMALLLEGNAHYRCDF<br>RTTYKAKEKGVKLPGAHFVDHCIEILSHDKDYNKVKLYEHA<br>VAHSGLPDNARRSGLRSRAQASNSAVDGTAGPGSTGSR |
| K249Q-hOGG1 | MPARALLPRRMGHRTLASTPALWASIPCPRSELRLDLVLPSG<br>QSFRWREQSPAHWSGVLADQVWTLTQTEEQLHCTVYRGDK<br>SQASRPTPDELEAVRKYFQLDVTLAQLYHHWGSVDSHFQEV<br>AQKFQGVRLLRQDPIECLFSFICSSNNNIARITGMVERLCQAF<br>GPRLIQLDDVTYHGFPSLQALAGPEVEAHLRKLGLGYRARY<br>VSASARAILEEQGGLAWLQQLRESSYEEAHKALCILPGVGT**Q**<br>VADCICLMALDKPQAVPVDVHMWHIAQRDYSWHPTTSQAK<br>GPSPQTNKELGNFFRSLWGPYAGWAQAVLFSADLRQSRHA<br>QEPPAKRRKGSKGPEG |
| H270A-hOGG1 | MPARALLPRRMGHRTLASTPALWASIPCPRSELRLDLVLPSG<br>QSFRWREQSPAHWSGVLADQVWTLTQTEEQLHCTVYRGDK<br>SQASRPTPDELEAVRKYFQLDVTLAQLYHHWGSVDSHFQEV<br>AQKFQGVRLLRQDPIECLFSFICSSNNNIARITGMVERLCQAF |

| | |
|---|---|
| | GPRLIQLDDVTYHGFPSLQALAGPEVEAHLRKLGLGYRARY<br>VSASARAILEEQGGLAWLQQLRESSYEEAHKALCILPGVGTK<br>VADCICLMALDKPQAVPVDV**<u>A</u>**MWHIAQRDYSWHPTTSQAK<br>GPSPQTNKELGNFFRSLWGPYAGWAQAVLFSADLRQSRHA<br>QEPPAKRRKGSKGPEG |
| F319A-hOG<br>G1 | MPARALLPRRMGHRTLASTPALWASIPCPRSELRLDLVLPSG<br>QSFRWREQSPAHWSGVLADQVWTLTQTEEQLHCTVYRGDK<br>SQASRPTPDELEAVRKYFQLDVTLAQLYHHWGSVDSHFQEV<br>AQKFQGVRLLRQDPIECLFSFICSSNNNIARITGMVERLCQAF<br>GPRLIQLDDVTYHGFPSLQALAGPEVEAHLRKLGLGYRARY<br>VSASARAILEEQGGLAWLQQLRESSYEEAHKALCILPGVGTK<br>VADCICLMALDKPQAVPVDVHMWHIAQRDYSWHPTTSQAE<br>GPSPQTNKELGNFFRSLWGPYAGWAQAVL**<u>A</u>**SADLRQSRHA<br>QEPPAKRRKGSKGPEG |
| 30TC | 5’-*TCTCTCTCTCTCTCTCTCTCTCTCTCTCTCTCTCTCTCTCTCT*<br>*CTCTCTCTCTCTCTCTC*-3’ |
| 30TC-OGMS<br>1 | 5-*TCTCTCTCTCTCTCTCTCTCTCTCTCTCTCTCTCTCTCTCTCT*<br>*CTCTCTCTCTCTCTCTC*TTTTTTTTTT<u>GTTTCGGACGACACC</u><br><u>GGTCAGCGGTAGCCTGTCGTCCCGAGA</u>-3’ |
| 30TC-OG10 | 5′-*TCTCTCTCTCTCTCTCTCTCTCTCTCTCTCTCTCTCTCTCTCT*<br>*CTCTCTCTCTCTCTC*TCTTTTTTTTTTT<u>CGGGACGACGGGCC</u><br><u>AGTAACTAGCTCACTCATTACACTTGGTCGTCCC</u>-3′ |
| 4GA-AF647 | 5′-GAGAGAGA-3′-AF647 |
| 4GA-ROX | 5′-GAGAGAGA-3′-ROX |
